# How Well Does Structural MRI Classify Autism? A Leakage-Free, Site-Aware and Externally Validated Machine-Learning Evaluation in Two ABIDE Cohorts

**DOI:** 10.64898/2026.10.31.763319

**Authors:** Barış Akar, Çağatay Elik, Tuğba Altıntaş

## Abstract

**Background:** Many studies report high accuracy for classifying autism spectrum disorder (ASD) from structural MRI, but small samples, single train–test splits, site effects and information leakage can inflate performance. We evaluated how well brain morphometry distinguishes autistic from typically developing (TD) individuals under rigorous, leakage-free validation.

**Methods:** FreeSurfer-derived features (n = 280: regional thickness, area and volume; subcortical volumes; Yeo-network composites; asymmetry indices; global measures) from ABIDE II (804 participants; 341 ASD; 12 sites) were used to train logistic regression, linear support vector machine, random forest, XGBoost and multilayer perceptron classifiers. Confound regression (age, age^2^, sex, brain volume), site-wise centring and standardisation were learned within training folds only. Performance was assessed with nested 10-fold cross-validation, leave-one-site-out validation and external validation in ABIDE I (917 participants; 20 sites), and compared with full-pipeline permutation tests and a confound-only baseline.

**Results:** Cross-validated discrimination was modest: the best model (regularised logistic regression on all features) reached an area under the ROC curve (AUC) of 0.60 [0.56, 0.64], significantly above chance (permutation *p* = .002) but far from clinically useful and below a confound-only baseline using age, sex, brain volume and site (AUC 0.63); non-linear models did not perform better (AUC 0.45–0.60 across models and feature sets). Pooled leave-one-site-out AUCs were 0.54–0.61, with large between-site variation, and AUCs in the external ABIDE I cohort were 0.52–0.58. Age-stratified models did not exceed AUC 0.60. Across 300 random 80/20 splits in young adults, test-set AUCs ranged from 0.21 to 0.90, showing that single-split estimates such as 0.74 can arise by chance.

**Conclusions:** Under leakage-free, site-aware and externally validated evaluation, structural MRI morphometry classifies autism only slightly better than chance. Reports of high accuracy should be interpreted with caution, and the heterogeneity of autism argues for stratification and normative approaches rather than binary diagnostic classifiers.

## 1. Introduction

Autism spectrum disorder (ASD) is diagnosed on the basis of behaviour, and the search for objective biological markers has motivated a large literature applying machine learning (ML) to brain MRI^1–3^. Early structural MRI studies reported classification accuracies of 80–90% in small samples^4^, and the open Autism Brain Imaging Data Exchange (ABIDE) made it possible to test such claims in more than 2,000 scans from over 30 sites^5,6^. In these larger, multi-site samples, performance has generally been much lower: anatomical features classified ASD with about 60% accuracy in ABIDE I^7^, sub-grouping by age, sex and IQ improved performance only for selected subgroups^8^, and functional connectivity approaches reached roughly 60–70% accuracy^9–11^. An international biomarker challenge on ABIDE-like data likewise found that predictions generalised only moderately^12^.

The gap between small-sample and large-sample results reflects well-known methodological pitfalls. Accuracy estimates from small samples are highly variable, and cross-validation error bars are large^13^. Hyperparameter tuning on the same data used for evaluation biases estimates upwards unless nested cross-validation is used^14^. Preprocessing steps such as harmonisation, confound removal, feature selection or scaling that are fitted on the full data before splitting leak information from the test set^15,16^. Scanner and site differences can be learned instead of biology, so that performance collapses in unseen sites, and evaluating models only on a single random train–test split ignores this variability^17,18^. Confounds such as age, sex and head size may also differ between groups and inflate apparent accuracy^16^.

A further consideration is the heterogeneity of ASD itself. Autistic individuals differ widely in aetiology, cognitive profile and brain anatomy, and group-level structural differences are small^19,20^. Normative modelling has shown that autistic individuals deviate from typical development in individually specific ways^21,22^; if so, binary classifiers would be expected to perform modestly regardless of algorithmic sophistication^23^.

We therefore evaluated, under conditions designed to avoid leakage and over-optimism, how well structural MRI morphometry classifies ASD. We compared five commonly used algorithms on three feature sets derived from FreeSurfer, learned all preprocessing within training folds, and assessed performance with nested cross-validation, leave-one-site-out validation and external validation in an independent cohort. We further quantified chance performance with full-pipeline permutation tests, compared against a confound-only baseline, examined age-stratified models and estimated the variability of single train–test splits, a design common in the literature.

## 2. Materials and Methods

### 2.1. Participants and data

We used FreeSurfer outputs and phenotypic data from ABIDE II^6^ (FreeSurfer 6.0, Mindboggle processing^24^) as the development cohort and ABIDE I^5^ (FreeSurfer 5.1, Preprocessed Connectomes Project) as the external validation cohort. Quality control and site selection followed our companion morphometric analysis: scans with excessive surface defects^25^, implausible brain-to-intracranial volume ratios, failed expert ratings or missing labels were excluded, and only sites with at least five participants per group were retained. The final development cohort comprised 804 participants (341 ASD, 463 TD; 12 sites; age 5.2–64.0 years) and the external cohort 917 participants (432 ASD, 485 TD; 20 sites). Sample characteristics are given in Table 1. All data are de-identified and publicly available; contributing sites obtained ethics approval and informed consent, and no additional ethics approval was required for this secondary analysis.

**Table 1.**
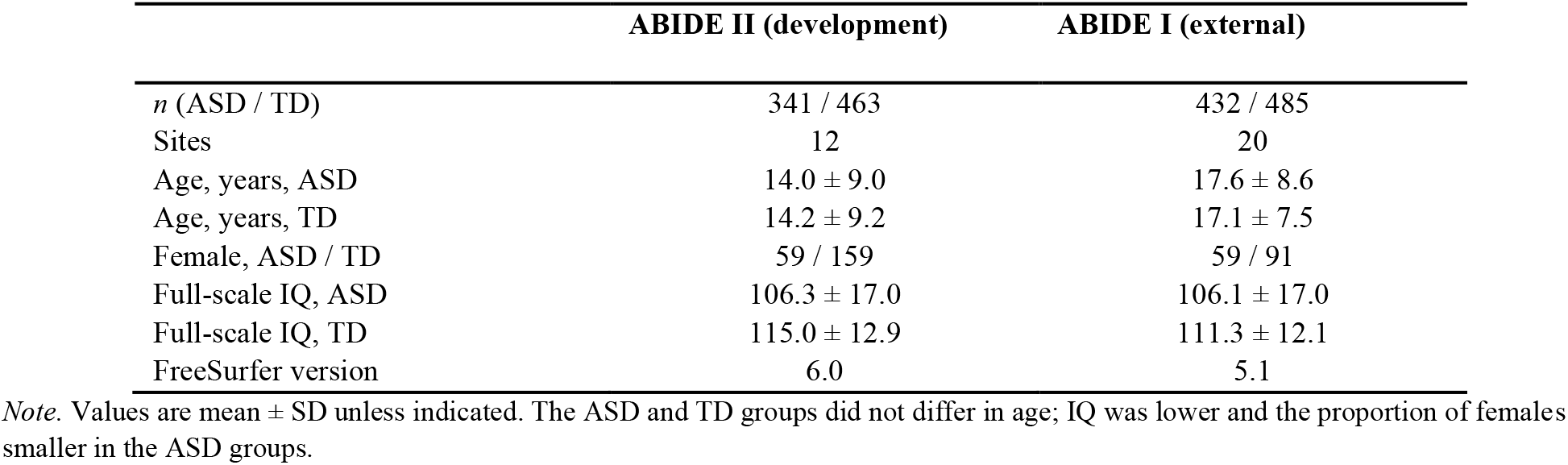
Characteristics of the Development (ABIDE II) and External Validation (ABIDE I) Cohorts.

### 2.2. Features

Three feature sets were defined from FreeSurfer outputs^26^. The *regional* set (110 features) contained the cortical thickness, surface area and grey matter volume of the 34 bilateral Desikan– Killiany regions^27^ and eight bilateral subcortical volumes. The *network* set (60 features) contained thickness, area and volume composite scores for the seven Yeo networks^28^ (regions weighted by their vertex overlap with each network), the corresponding network asymmetry indices, hemispheric asymmetry indices and global measures (cortical and subcortical grey matter, white matter, cerebellum, corpus callosum and its segments, ventricles, mean thickness and total area). The *all* set (280 features) combined both sets with regional and subcortical asymmetry indices, AI = 100 × (L − R)/[(L + R)/2].

### 2.3. Leakage-free preprocessing

All preprocessing parameters were estimated on training data only and then applied to test data. Within each training fold, each feature was regressed on age, age^2^, sex and (for volumes and areas) total brain volume, and the resulting coefficients were used to residualise training and test data. Because site effects cannot be estimated for unseen sites with supervised harmonisation without using diagnostic labels of the test site^29,30^, residuals were centred on their site median within each set; this label-free operation uses no diagnostic information. Features were then standardised with training-set means and standard deviations. Total brain volume was used instead of intracranial volume because intracranial volume estimates were unreliable in the ABIDE I release.

### 2.4. Classifiers and validation

Five classifiers were compared: L2-regularised logistic regression and linear support vector machine (SVM) with class-balanced weights, random forest^31^ (500 trees), gradient-boosted trees (XGBoost^32^; 300 trees) and a multilayer perceptron (MLP) with early stopping. Hyperparameters (regularisation strength; tree depth, minimum leaf size and learning rate; hidden-layer size and weight decay) were selected by an inner stratified 5-fold grid search maximising the area under the receiver operating characteristic curve (AUC) within each outer training fold (nested cross-validation^14^).

Performance was evaluated in three ways. (i) Nested stratified 10-fold cross-validation in ABIDE II, with folds stratified by site and diagnosis; out-of-fold predictions were pooled to compute AUC with 95% bootstrap confidence intervals (2,000 resamples), balanced accuracy, sensitivity and specificity. (ii) Leave-one-site-out (LOSO) validation, in which each ABIDE II site was held out in turn, using the all-feature set. (iii) External validation, in which models trained on all ABIDE II participants were applied unchanged to ABIDE I. Reporting follows TRIPOD recommendations where applicable^33^.

### 2.5. Reference analyses

Chance performance was estimated with full-pipeline permutation tests^34^: diagnostic labels were permuted and the entire cross-validation procedure, including preprocessing, was repeated (500 permutations for logistic regression with fixed hyperparameters). A confound-only baseline used age, age^2^, sex, total brain volume and site indicators as the only predictors. Age-stratified models were trained separately in childhood (< 12 years), adolescence (12–17 years) and adulthood (≥ 18 years). To quantify the variability of the single-split evaluations common in the literature, we repeated an 80/20 stratified train–test split 300 times in participants aged 19–30 years using XGBoost with leakage-free preprocessing. Model explanations were obtained from the regularised logistic regression coefficients and from SHAP values of an XGBoost model fitted to all ABIDE II data^35^.

### 2.6. Use of generative AI tools

In line with publisher policy, the authors declare that a large language model–based assistant (Claude, Anthropic) was used to help write the analysis code, produce figures and draft and edit the manuscript text. All analysis decisions, outputs, references (each verified against Crossref records) and the final text were checked by the authors, who take full responsibility for the content. No AI tool is listed as an author.

## 3. Results

### 3.1. Cross-validated performance

Cross-validated performance was modest for every model and feature set (Table 2, Figure 2a, b). The highest AUC was obtained by logistic regression on the all-feature set (0.60 [0.56, 0.64]; balanced accuracy 0.57), closely followed by the linear SVM on all features (0.59 [0.55, 0.63]) and logistic regression on regional features (0.59 [0.55, 0.63]). Non-linear models did not outperform linear ones: random forest, XGBoost and MLP reached AUCs of 0.45–0.57, and the MLP performed at chance on the regional and network sets. Network composite features alone performed worse than regional features with linear models. Strikingly, the confound-only baseline—age, age^2^, sex, brain volume and site, without any regional brain feature—reached AUC = 0.63 [0.60, 0.67], higher than every brain-based model. Because diagnostic proportions and sex ratios differed between sites, sample composition alone carried diagnostic information; a model that does not remove these confounds could therefore appear to perform well without learning anything about brain anatomy.

**Table 2.** Nested 10-Fold Cross-Validated Classification Performance in ABIDE II.

| **Feature set** | **Model** | **AUC [95% CI]** | **Balanced accuracy** | **Sensitivity** | **Specificity** |
|---|---|---|---|---|---|
| Regional | Logistic regression | 0.59 [0.55, 0.63] | 0.58 | 0.53 | 0.62 |
| Regional | Linear SVM | 0.59 [0.55, 0.63] | 0.57 | 0.51 | 0.63 |
| Regional | Random forest | 0.57 [0.53, 0.61] | 0.54 | 0.24 | 0.84 |
| Regional | XGBoost | 0.56 [0.52, 0.60] | 0.55 | 0.34 | 0.77 |
| Regional | MLP | 0.46 [0.42, 0.50] | 0.49 | 0.13 | 0.86 |
| Network | Logistic regression | 0.54 [0.50, 0.58] | 0.53 | 0.54 | 0.52 |
| Network | Linear SVM | 0.54 [0.49, 0.57] | 0.51 | 0.49 | 0.54 |
| Network | Random forest | 0.57 [0.53, 0.61] | 0.53 | 0.25 | 0.81 |
| Network | XGBoost | 0.57 [0.53, 0.61] | 0.55 | 0.35 | 0.75 |
| Network | MLP | 0.45 [0.41, 0.49] | 0.47 | 0.28 | 0.65 |
| All | Logistic regression | 0.60 [0.56, 0.64] | 0.57 | 0.56 | 0.57 |
| All | Linear SVM | 0.59 [0.55, 0.63] | 0.57 | 0.57 | 0.57 |
| All | Random forest | 0.56 [0.52, 0.60] | 0.54 | 0.21 | 0.87 |
| All | XGBoost | 0.55 [0.51, 0.60] | 0.54 | 0.28 | 0.79 |
| All | MLP | 0.55 [0.51, 0.59] | 0.55 | 0.45 | 0.65 |

*Note. n* = 804. AUC confidence intervals from 2,000 bootstrap resamples of out-of-fold predictions. Regional: 110 features; network: 60 features; all: 280 features. Sensitivity and specificity at a probability threshold of 0.5 (decision value 0 for SVM).

**Figure 1.**
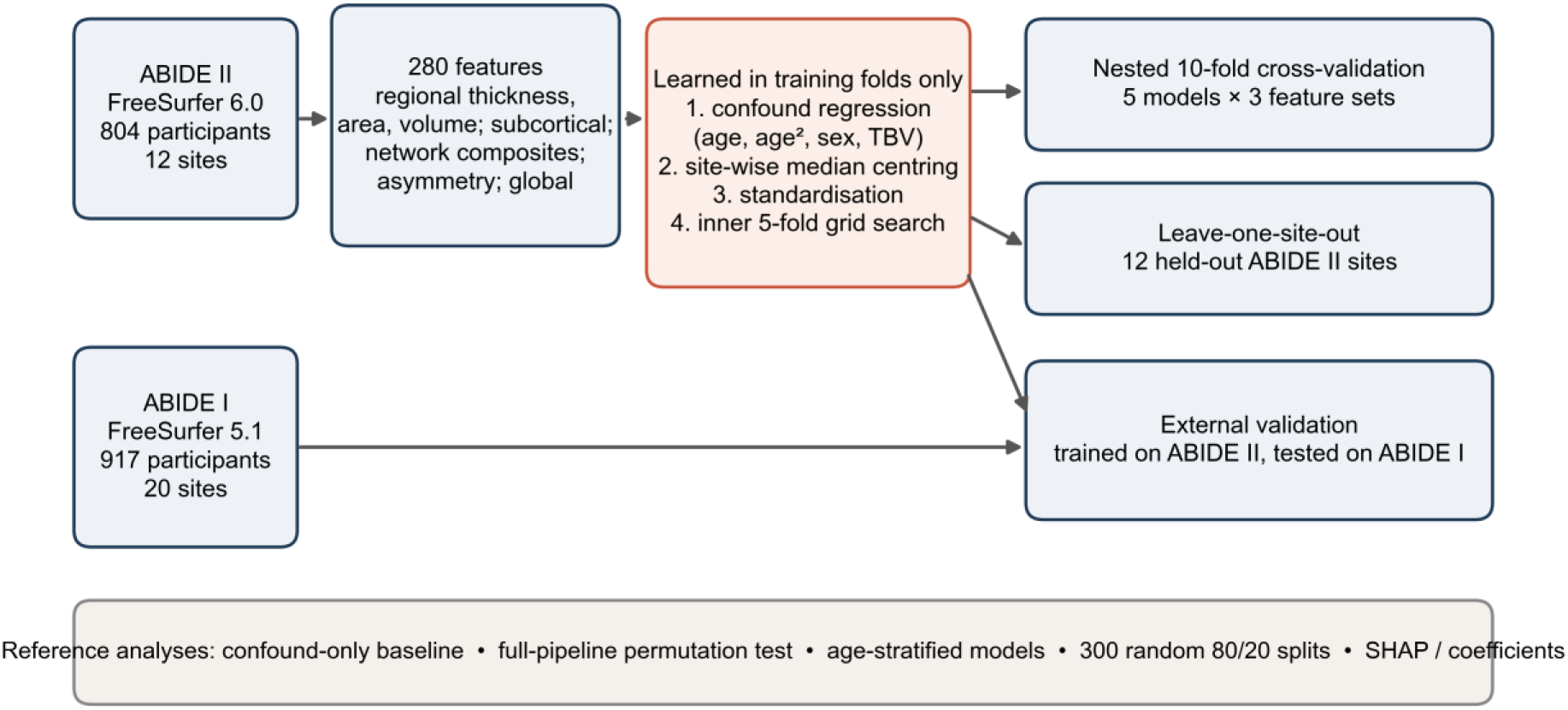
Analysis pipeline. All preprocessing (confound regression, site-wise centring, standardisation) and hyperparameter selection were learned within training data only. Performance was assessed with nested 10-fold cross-validation, leave-one-site-out validation and external validation in ABIDE I, and compared with permutation-based chance levels, a confound-only baseline, age-stratified models and the variability of single train–test splits.

**Figure 2.**
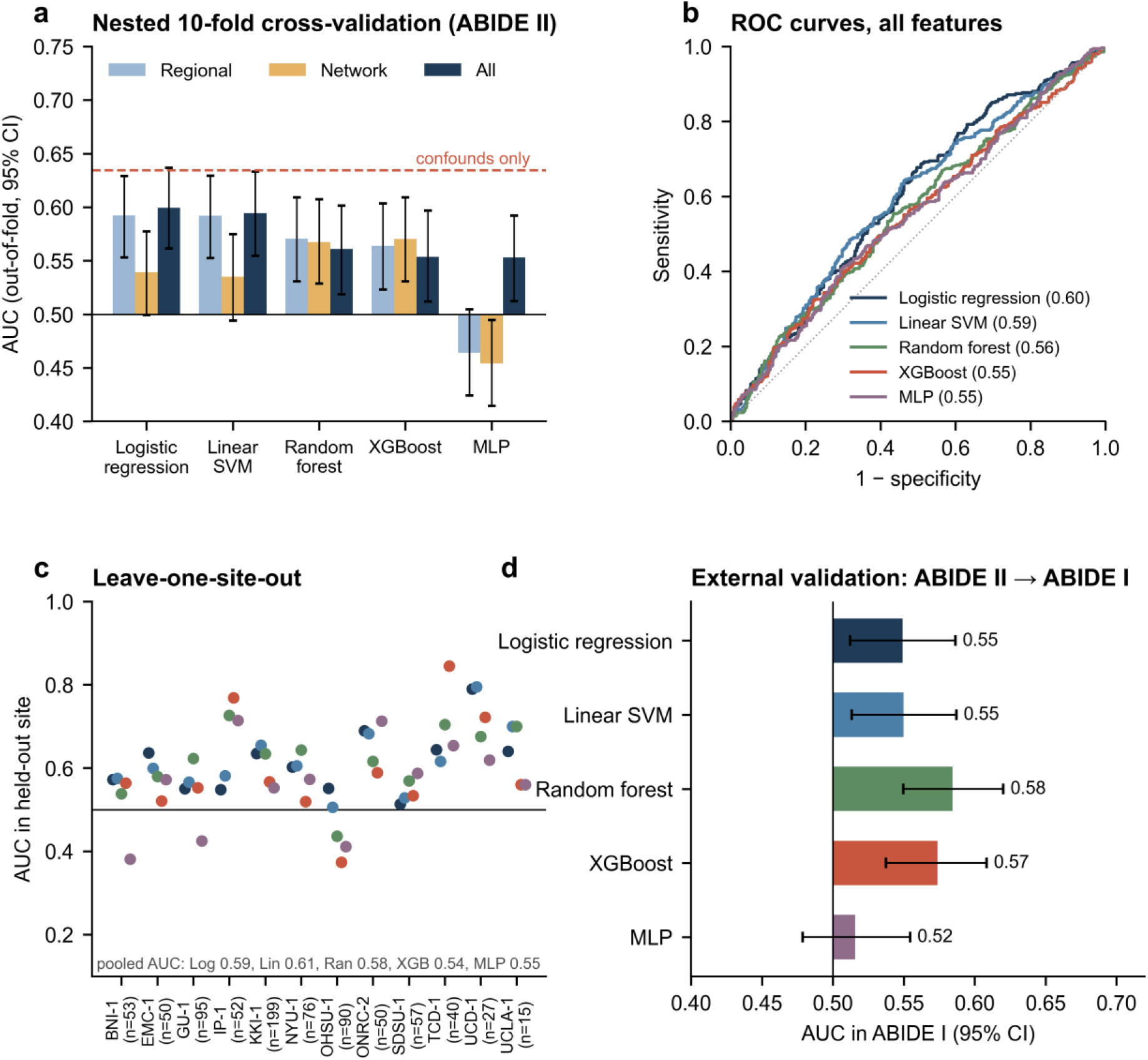
Classification performance. (a) Out-of-fold AUC (95% bootstrap CI) of five classifiers on three feature sets in nested 10-fold cross-validation (ABIDE II); the dashed line shows the confound-only baseline. (b) ROC curves for the all-feature set. (c) AUC in each held-out site in leave-one-site-out validation (*n* per site in parentheses). (d) External validation: models trained on ABIDE II and tested on ABIDE I.

Despite its modest size, performance exceeded chance: in the full-pipeline permutation test, logistic regression with fixed hyperparameters reached AUC = 0.61, compared with a null mean of 0.50 and a 95th percentile of 0.54 (*p* = .002; Figure 3a).

**Figure 3.**
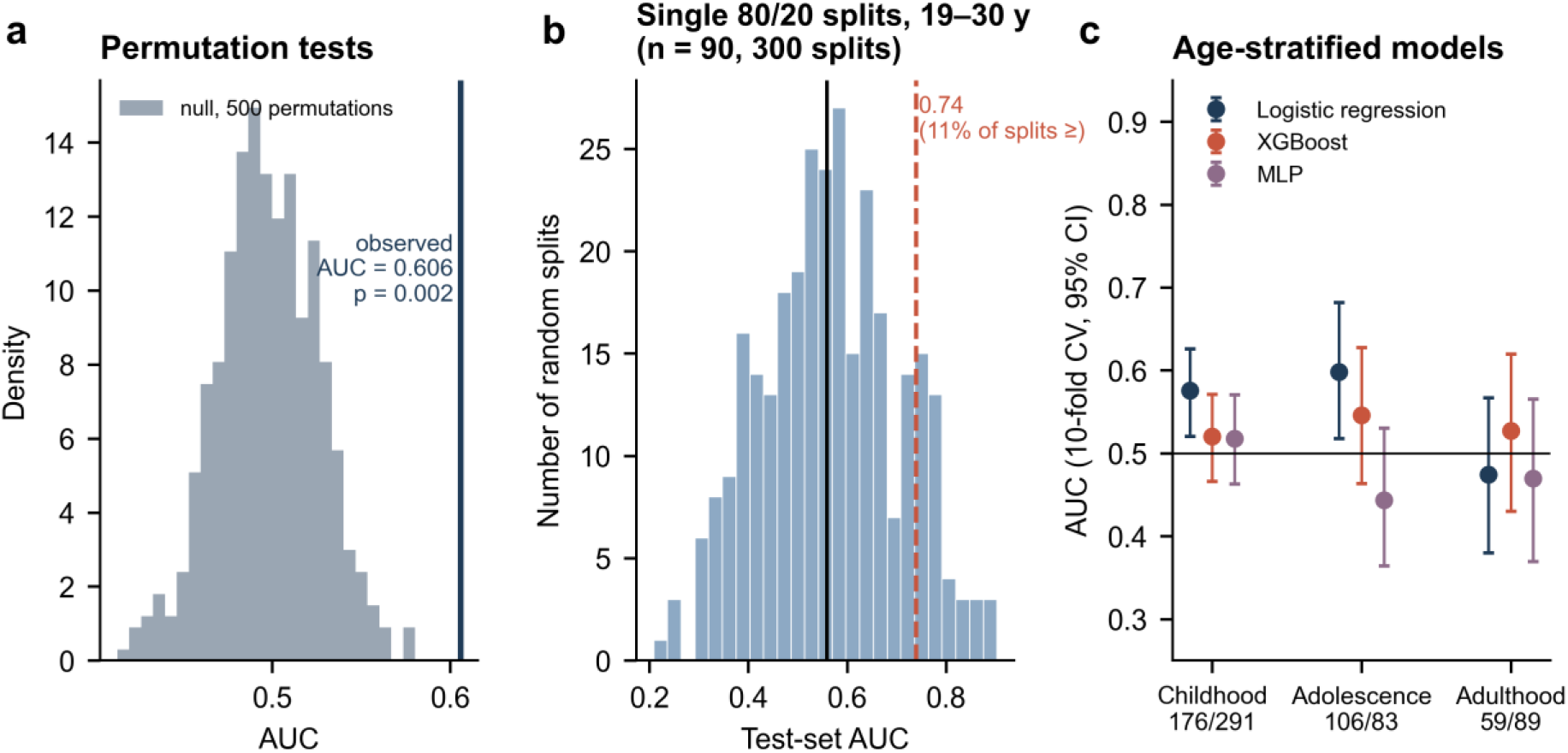
Reference analyses. (a) Full-pipeline permutation null distribution and observed AUC (vertical line). (b) Distribution of test-set AUCs across 300 random 80/20 train–test splits in participants aged 19–30 years; the dashed line marks AUC = 0.74 and the solid line the mean. (c) Age-stratified cross-validated AUCs (95% CI) with group sizes (ASD/TD).

### 3.2. Generalisation to new sites and an independent cohort

When entire sites were held out, pooled AUCs ranged from 0.54 (XGBoost) to 0.61 (Linear SVM), similar to cross-validation, but performance varied widely across held-out sites (site-level AUC 0.37–0.84; Figure 2c). In the external ABIDE I cohort, processed with a different FreeSurfer version and collected at different sites, AUCs were 0.52–0.58; the best external performance was obtained by random forest (0.58 [0.55, 0.62]; Figure 2d, Table 3).

**Table 3.**
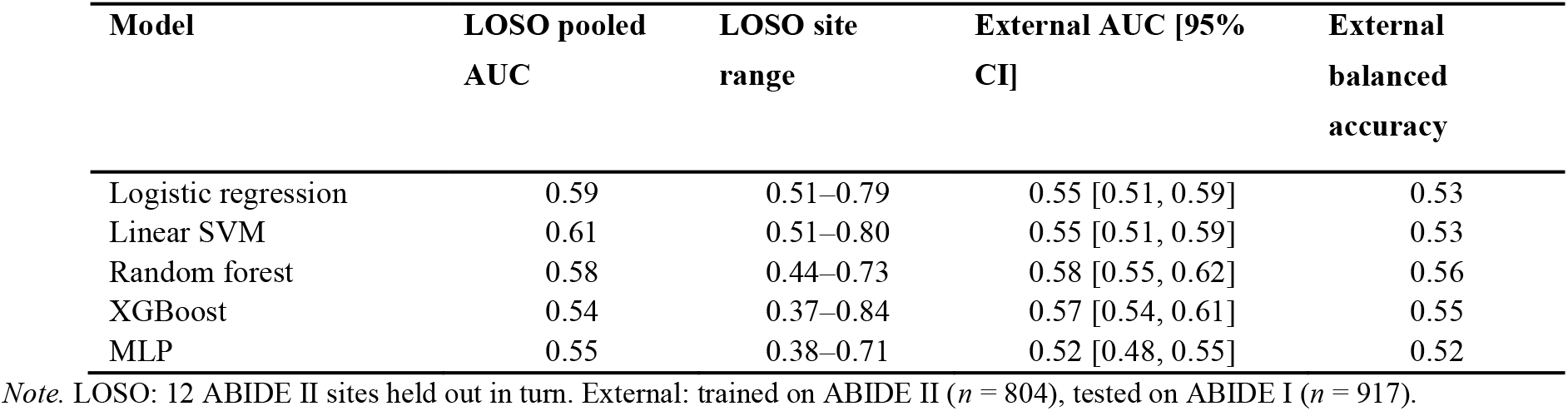
Leave-One-Site-Out and External Validation (All-Feature Set)

### 3.3. Age-stratified models and single-split variability

Training separate models within developmental stages did not improve performance (Figure 3c). The best stage-specific AUCs were 0.58 in childhood, 0.60 in adolescence and 0.53 in adulthood, with wide confidence intervals in the smaller strata. Across 300 random 80/20 splits in participants aged 19–30 years (*n* = 90), test-set AUCs of the same XGBoost pipeline ranged from 0.21 to 0.90 (mean 0.56), and 11% of splits produced AUC ≥ 0.74 (Figure 3b). Thus, a single split can yield apparently good performance by chance alone.

### 3.4. Model explanations

The features contributing most to the XGBoost model (mean |SHAP|) were posterior cingulate thickness, superior temporal thickness, precentral volume, subcortical grey matter and Cerebral white matter hemispheric asymmetry (Figure 4a). The largest logistic regression coefficients involved posterior cingulate thickness (higher in ASD); banks of the superior temporal sulcus thickness asymmetry (higher in ASD); paracentral surface area (higher in ASD); lateral orbitofrontal thickness (lower in ASD) (Figure 4b; for asymmetry indices, higher values indicate more leftward asymmetry). Individual feature contributions were small and distributed across many regions, consistent with the absence of a single dominant anatomical marker.

**Figure 4.**
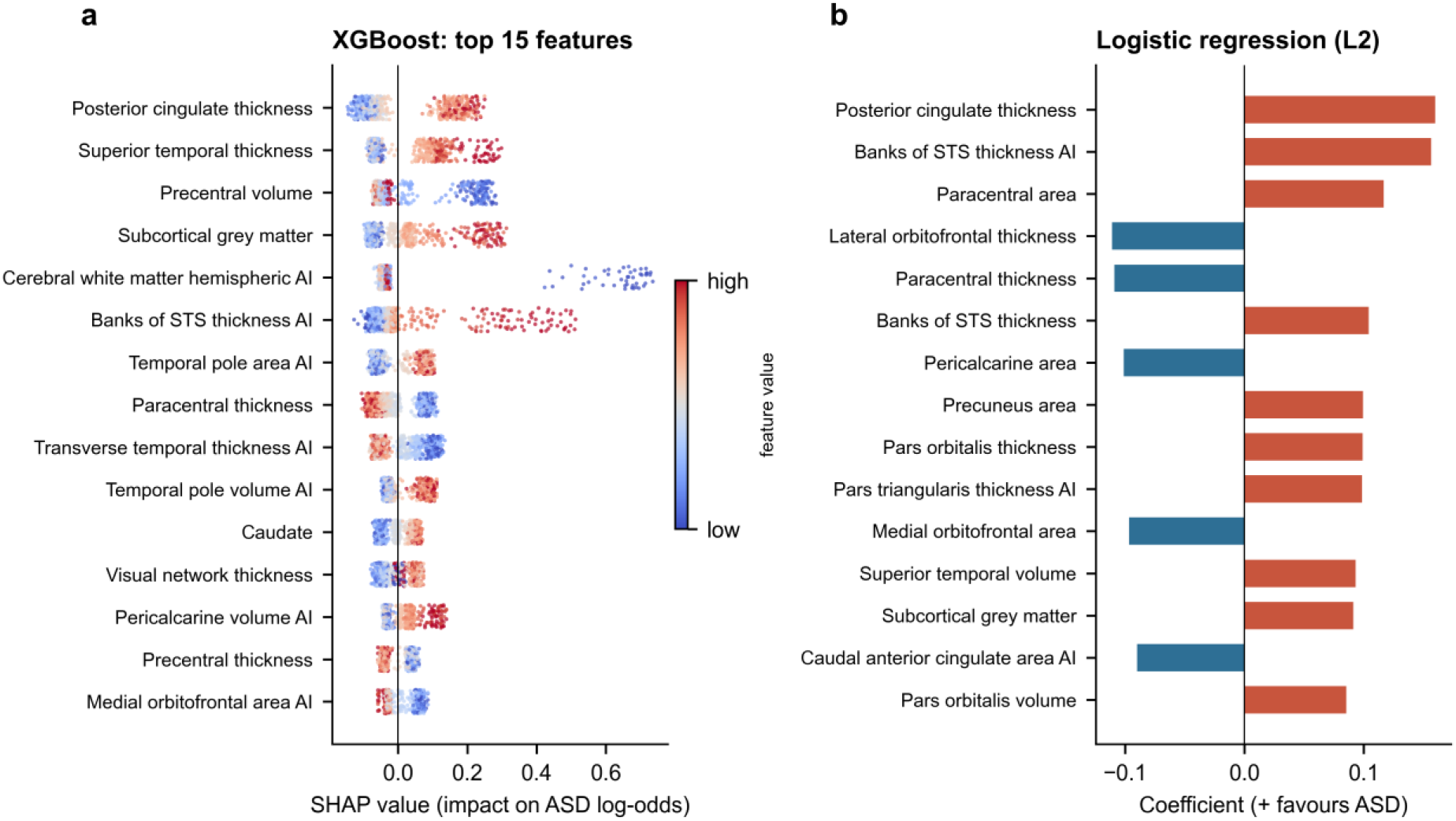
Model explanations. (a) SHAP values of the 15 most important features of an XGBoost model trained on all ABIDE II data; each point is one participant, coloured by feature value. (b) The 15 largest standardised coefficients of the L2-regularised logistic regression (positive: higher values predict ASD). AI, asymmetry index.

## 4. Discussion

Using leakage-free preprocessing, nested cross-validation, leave-one-site-out and external validation in two independent cohorts, we found that structural MRI morphometry distinguishes autistic from typically developing individuals only modestly. The best models reached AUCs of about 0.60, significantly above chance but far below the level required for clinical use, and more flexible algorithms offered no advantage over regularised linear models. Performance fell further in an external cohort and varied widely between held-out sites.

These results agree with large-sample ABIDE studies reporting accuracies around 60% for anatomical features^7,8^ and with the modest generalisation observed in a biomarker challenge^12^, and they contrast with the high accuracies of early small-sample studies^4^. Our reference analyses illustrate why such discrepancies arise. First, a model using only age, sex, brain volume and site outperformed all brain-based models, because group composition differed between sites; without in-fold confound removal, such sampling artefacts are easily mistaken for neurobiological signal. Second, single train– test splits produced a wide range of AUCs from the same data and pipeline, so that an apparently strong result (e.g., AUC 0.74) can occur by chance; small test sets yield large error bars^13^. Tuning hyperparameters without nesting^14^, fitting harmonisation or scaling on the full data^15^ and ignoring confounds^16^ further inflate estimates. Best-practice recommendations—external validation, transparent reporting and appropriate baselines^17,18,33^—are therefore essential for neuroimaging biomarkers in psychiatry.

The modest performance is also consistent with the biology of autism. Group differences in morphometry are small^19^, and autistic individuals deviate from normative expectations in individually specific ways^22^; our companion analysis of the same data found more frequent but spatially idiosyncratic extreme deviations in autism. If anatomical alterations differ from person to person, a single decision boundary cannot separate the groups well, no matter how flexible the classifier^20,23^. Model explanations supported this view: predictive information was spread thinly across many regions, including visual and temporal cortex previously implicated in autism^36,37^, rather than concentrated in a few markers. Future work may therefore gain more from stratifying autistic individuals into biologically coherent subgroups, from normative deviation features^21^ and from multimodal data than from more complex classifiers applied to case–control labels.

### 4.1. Limitations

Our results are limited to FreeSurfer morphometry summarised in an atlas; vertex-wise or deep-learning approaches applied to raw images might capture additional information, although previous large studies suggest modest gains^11^. The development and external cohorts differed in FreeSurfer version, which may have reduced external performance. Label-free site centring removes site differences without using diagnostic labels but may also remove genuine group differences in sites with unbalanced diagnosis ratios. Diagnostic procedures, IQ and sex ratios varied across sites, and adults, females and individuals with intellectual disability were under-represented. Finally, permutation testing with all five models was computationally prohibitive, so it was performed for the linear model with fixed hyperparameters.

## 5. Conclusions

Under rigorous, leakage-free and externally validated evaluation, structural MRI morphometry classifies autism only slightly better than chance, and sophisticated non-linear models do not change this conclusion. Claims of high diagnostic accuracy from structural MRI should be treated with caution, and research efforts may be better directed towards understanding the heterogeneity of autism than towards binary classification.

## Declarations

### Ethics

Secondary analysis of de-identified, publicly available ABIDE data; all contributing sites obtained ethics approval and informed consent, and no additional ethics approval was required.

### Data availability

ABIDE data are openly available (http://fcon_1000.projects.nitrc.org/indi/abide/); FreeSurfer outputs were obtained from the public fcp-indi repository. Derived features and analysis code are available from the corresponding author upon reasonable request.

### Author contributions

B.A.: conceptualisation, data curation, analysis, writing—original draft. Ç.E.: methodology (machine learning), writing—review and editing. T.A.: supervision, methodology, writing—review and editing. All authors approved the final manuscript.

## Acknowledgements

We thank the ABIDE consortium, its contributing sites and participants, and the Preprocessed Connectomes Project.

